# Multi-Layered Maps of Neuropil with Segmentation-Guided Contrastive Learning

**DOI:** 10.1101/2022.03.29.486320

**Authors:** Sven Dorkenwald, Peter H. Li, Michał Januszewski, Daniel R. Berger, Jeremy Maitin-Shepard, Agnes L. Bodor, Forrest Collman, Casey M. Schneider-Mizell, Nuno Maçarico da Costa, Jeff W. Lichtman, Viren Jain

**Author notes:** co-first authors (Sven Dorkenwald, Peter H. Li).

## Abstract

Maps of the nervous system that identify individual cells along with their type, subcellular components, and connectivity have the potential to reveal fundamental organizational principles of neural circuits. Volumetric nanometer-resolution imaging of brain tissue provides the raw data needed to build such maps, but inferring all the relevant cellular and subcellular annotation layers is challenging. Here, we present Segmentation-Guided Contrastive Learning of Representations (“SegCLR”), a self-supervised machine learning technique that produces highly informative representations of cells directly from 3d electron microscope imagery and segmentations. When applied to volumes of human and mouse cerebral cortex, SegCLR enabled the classification of cellular subcompartments (axon, dendrite, soma, astrocytic process) with 4,000-fold less labeled data compared to fully supervised approaches. Surprisingly, SegCLR also enabled inference of cell types (neurons, glia, and subtypes of each) from fragments with lengths as small as 10 micrometers, a task that can be difficult for humans to perform and whose feasibility greatly enhances the utility of imaging portions of brains in which many neuron fragments terminate at a volume boundary. These predictions were further augmented via Gaussian process uncertainty estimation to enable analyses restricted to high confidence subsets of the data. Finally, SegCLR enabled detailed exploration of layer-5 pyramidal cell subtypes and automated large-scale statistical analysis of upstream and downstream synaptic partners in mouse visual cortex.

## Introduction

Biological understanding has been enabled by annotating parts of organisms and elucidating their interrelationships. In the brain, numerous types of neuronal and glial cells have been discovered and cataloged according to their morphological, physiological, and molecular properties^1–5^, typically using methods that interrogate cells in a sparse or isolated setting. Further discoveries would be enabled by producing maps that contain dense assemblies of cells and multiple layers of annotation in the context of a neural circuit or region^6–10^.

Producing dense maps of neuropil is challenging due to the multiple scales of brain structures (e.g., nanometers for a synapse versus millimeters for an axon)^11^, and the vast number of objects within neuropil which must be individually segmented, typed, and annotated with their interrelationships. Volumetric electron microscopy (EM) has proven to be an effective way to image brain structures over both large and small scales^12–14^ and automated segmentation of volume EM data has also shown significant progress^15–19^, including the demonstration of millimeter-scale error-free run lengths^20^.

Here, we address the problem of efficiently inferring types and annotations of segmented structures by introducing Segmentation-Guided Contrastive Learning of Representations (SegCLR), a machine learning approach that is scalable in three important respects: first, the representations produced by a single SegCLR model can be used for a diverse set of annotation tasks (e.g., local identification of cellular subcompartments, assigning type to an entire cell or fragment); second, the compact representations learned by SegCLR enable accurate downstream analyses with simple linear classifiers or shallow networks, removing the need to train large models (such as 3d convolutional networks) for each additional task; and third, SegCLR reduces the amount of ground truth labeling required for specific tasks by up to 4 orders of magnitude. Perhaps most intriguingly, we show that SegCLR enables a type of annotation which is challenging for either automated methods or human experts: inferring the cell type from a short length (∼10-50 μm) of cortical cell fragment. This capability has important implications for the utility of cortical EM datasets that so far encompass only subsets of whole brains.

Previous machine learning methods for neuropil annotation have primarily used features that were hand-designed or derived from supervised learning, including random forests trained on hand-designed features^9,21^, 2d convolutional networks trained on projections of neuropil (“Cellular Morphology Networks”)^22,23^, point cloud networks trained on representations of cell membranes^24^, or 3d convolutional networks trained directly on voxels^25^. Schubert et al.^22,24^ trained cell representations using a triplet loss, but it was not reported whether these representations are suitable for downstream analyses. Previous results on cell type classification of neurite fragments required larger spatial context, precomputed organelle masks, and manually engineered features^21–24^, or used a single local view to achieve modest classification accuracy on a limited set of classes^22^.

Self-supervised learning has emerged as a broadly successful technique for producing representations of text^26^ and images^27^ without the use of labeled data. Weis *et al*. introduced a method for self-supervised learning of neuronal morphology that operates on coarse skeleton representations of neurons^28^; however, similar to previous morphological clustering methods^29–31^, this approach did not demonstrate important capabilities of SegCLR (e.g. classification of subcellular structures and type inference from small cellular fragments). Self-supervised learning has also been previously explored for content-based image retrieval in unsegmented 3d connectomic datasets^32^, and for image retrieval and classification in 2d medical image datasets^33^.

SegCLR takes inspiration from recent advances in self-supervised contrastive learning^27^ while introducing a segmentation-guided loss function in which positive example pairs are drawn from nearby, but not necessarily overlapping, cutouts of the same segmented cell. In addition to raw volumetric data, this approach therefore also requires 3d segmentation of individual cells throughout the volume, which is a typical requirement for subcellular annotation methods. As we demonstrate, current automated segmentation methods are sufficiently accurate to be used to train SegCLR without further human proofreading.

In addition to pursuing accurate classifications, it is critical to be able to estimate the uncertainty associated with specific predictions, such that biological inferences can be restricted to high confidence results. We show that SegCLR can be combined with Gaussian processes^34^ to provide a highly practical means of uncertainty estimation. Finally, we demonstrate the application of SegCLR to novel biological inference by detailed cell type analysis of upstream and downstream synaptic partners in mouse visual cortex.

## Results

### Training and inference of SegCLR embeddings

Representations that track separate cells through dense neuropil, such as instance segmentations^20^ or skeletonizations^35^, have proven fundamental to biological interpretation, as have annotations of select features of interest such as synapses^21,36^, organelles^37^, and cellular subcompartments^25^. To improve these methods, SegCLR produces tractable “embeddings”: vector representations that capture rich biological features in a dimensionally reduced space, and in which vector distance maps to a concept of biological distinctness (Fig. 1). These embeddings capture features relevant to a range of downstream tasks, and can be trained without manual feature engineering. Depending on the downstream application, embeddings can also be deployed without any requirement for manual proofreading or ground truth labeling, or with these requirements significantly reduced^38^. Each SegCLR embedding represents a local 3d view of EM data, and is focused on an individual cell or cell fragment within dense neuropil via an accompanying segmentation. Computed for billions of local views across large connectomic datasets, embeddings can directly support local annotation tasks (Fig. 2), or be flexibly combined at larger scales to support annotation at the level of cells and cell fragments (Figs. 3-5), or local circuits (Fig. 6).

**Figure 1.**
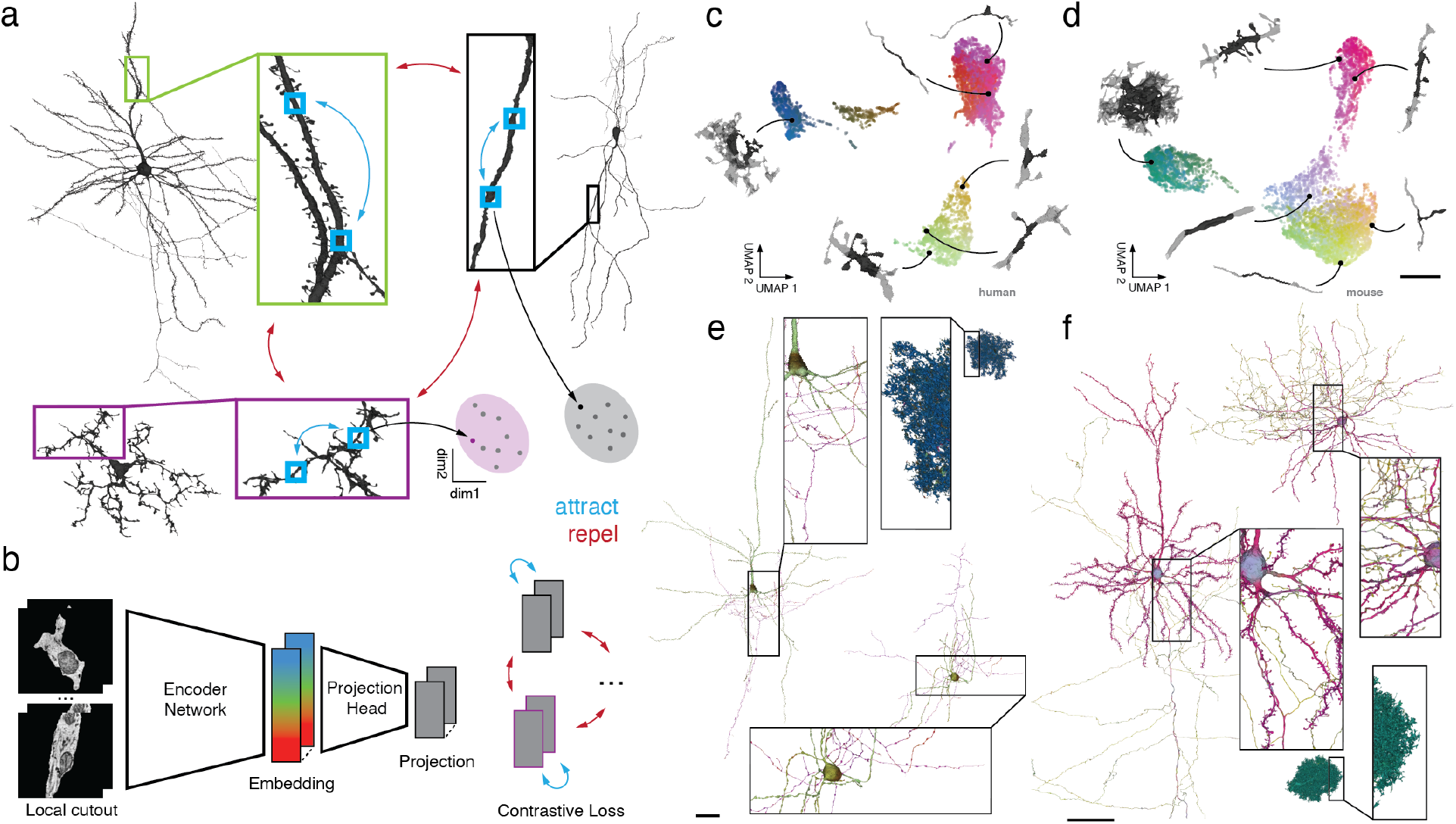
SegCLR: Segmentation-Guided Contrastive Learning of Representations. **a**. In SegCLR, positive pairs (blue double-headed arrows) are chosen from proximal but not necessarily overlapping 3d views (small blue boxes) of the same segmented cell, while negative pairs (red double-headed arrows) are chosen from different cells. The SegCLR network is trained to produce an embedding vector for each local 3d view such that embeddings are more similar for positive pairs compared to negative pairs (cartoon of clustered points). **b**. The input to the embedding network is a local 3d view (4.1×4.1×4.3 μm at 32×32×33 nm resolution for human data; 4.1×4.1×5.2 μm at 32×32×40 nm resolution for mouse) from the EM volume, masked by the segmentation for the object at the center of the field of view. A encoder network based on a ResNet-18 is trained to produce embeddings, via projection heads and a contrastive loss that are used only during training. **c-d**. Visualization via UMAP projection of the SegCLR embedding space for the human temporal cortex and mouse visual cortex datasets. Points for a representative sample of embeddings are shown, colored via 3d UMAP RGB, with the corresponding 3d morphology illustrated for 6 locations (network input in black, surrounded by 10×10×10 μm context in gray). Biologically related objects cluster naturally in this space without any ground truth label supervision, e.g. myelinated axons and axon initial segments occupy a space adjacent to the rest of the axons in the mouse embedding space. Scale bars (c,d): 5 μm **e-f**. Embeddings visualized along the extent of representative human (e) and mouse (f) cells. Each mesh rendering is colored according to the 3d UMAP RGB of the nearest embedding for the surrounding local 3d view. The embedding colors readily distinguish different types of cells, as well as different subcellular structures. Some axons are cut off to fit. Scale bars (e,f): 100 μm.

**Figure 2.**
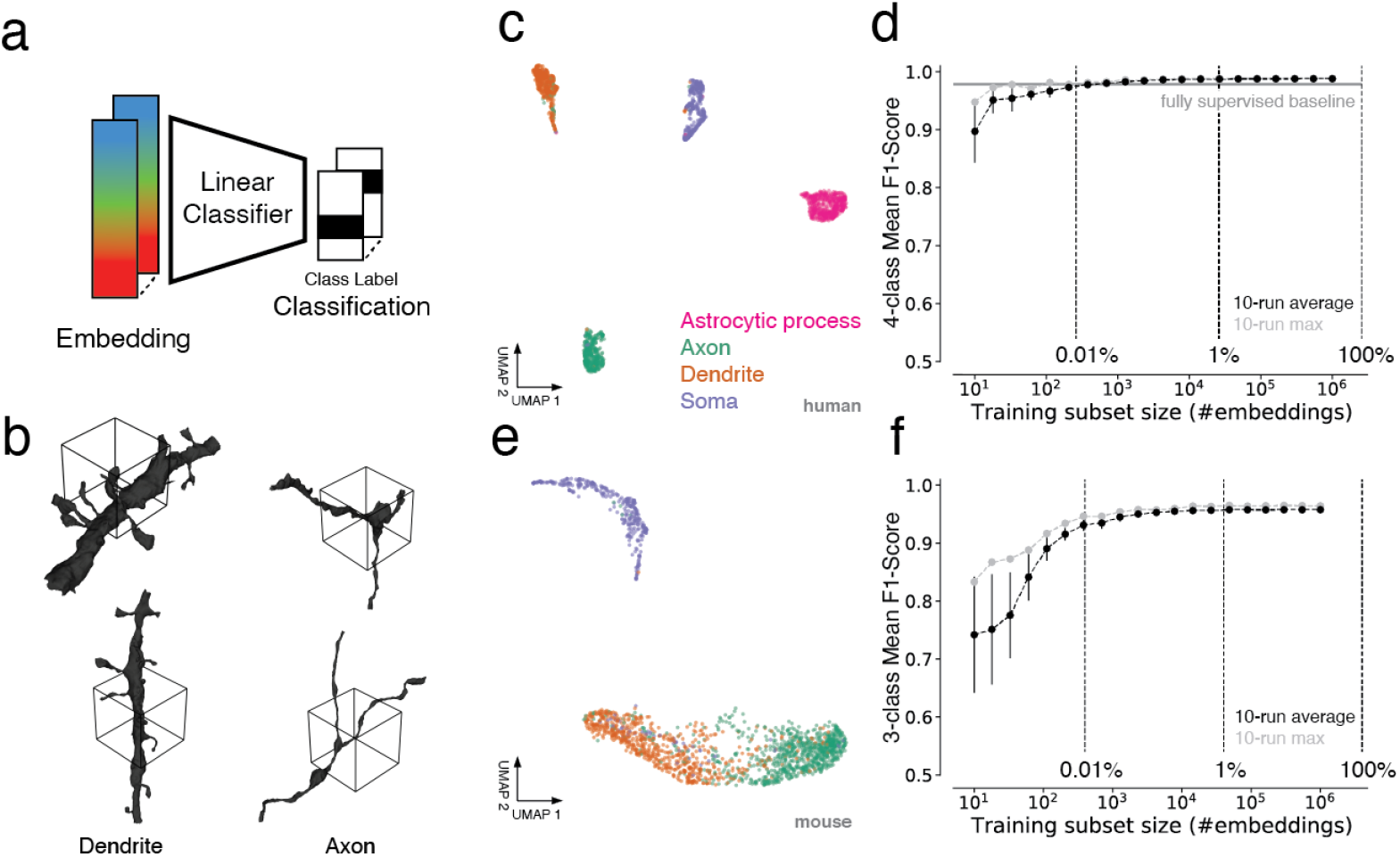
Subcompartment classification of SegCLR embeddings. **a**. Embedding vectors computed across the extent of the EM datasets can be used as compact inputs to downstream tasks, such as subcompartment classification. Each embedding represents a single local view (∼4-5 μm on a side). **b**. Ground truth examples of axon and dendrite subcompartment classes from the human temporal cortex dataset. The local 3d views for single embeddings is indicated by the wireframe cubes. The embedding network also receives the EM image data from within the segment mask (not shown). **c**. Embedding clusters from the human cortical dataset visualized via 2d UMAP. Each point is an embedding, colored by its ground truth subcompartment class as judged without reference to the embeddings. **d**. Evaluation of linear classifiers trained for the subcompartment task on the human dataset. The mean F1 score across classes was computed for networks trained using varying-sized subsets of the full available training data. For each training set sample size, mean and standard deviation of multiple subset resamplings is shown (error bars are obscured by the points for larger sample sizes). Light gray points show the best class-wise mean F1 score obtained for any training subset sampled at a given size. The horizontal line indicates the performance of a fully supervised ResNet-18 classifier trained on the full available training data. **e**. As in (c), for the mouse visual cortex dataset and three ground truth classes (axon, dendrite, soma). **f**. As in (d), for the mouse dataset.

**Figure 3.**
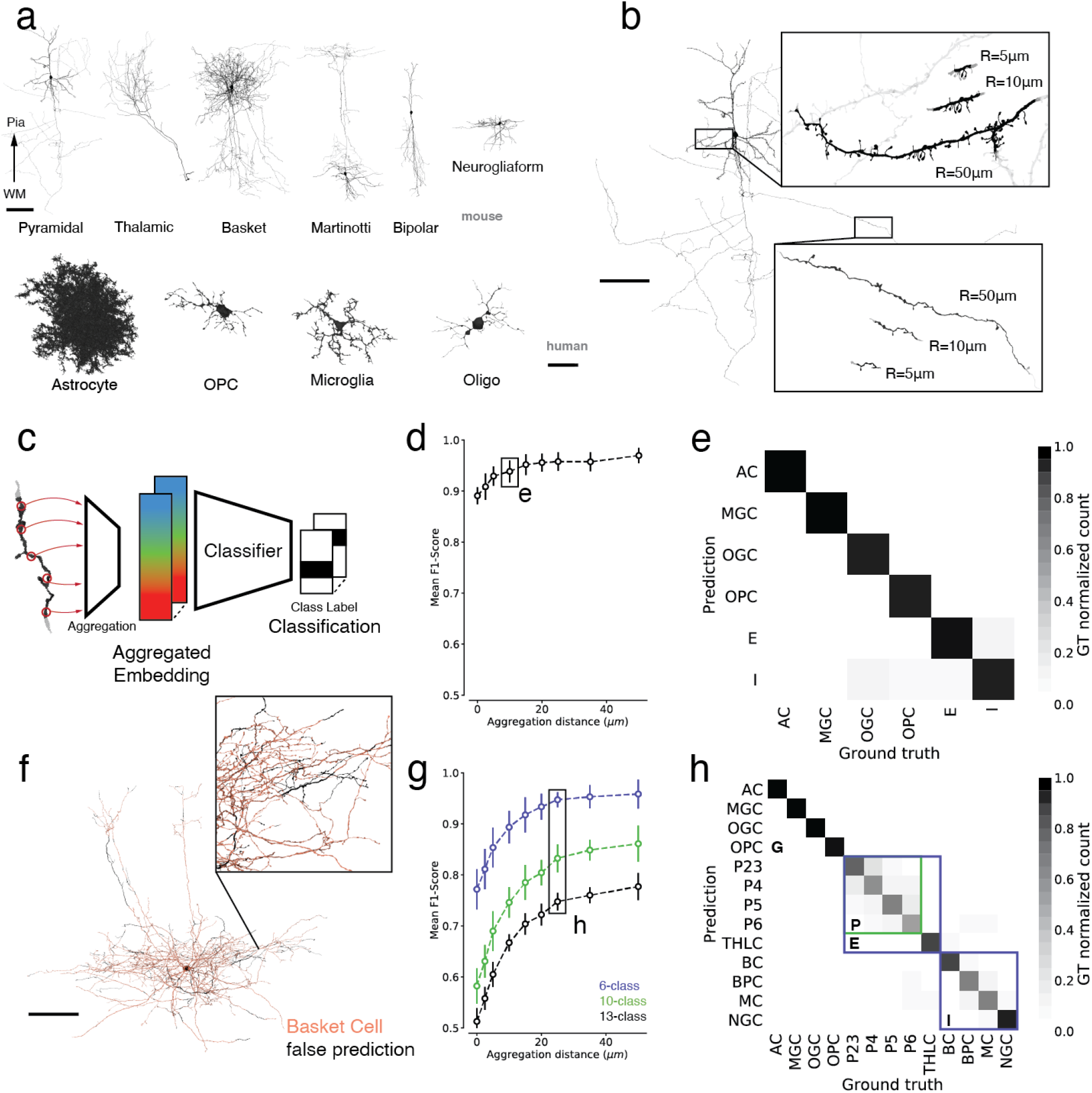
Cell type classification of large and small cell fragments via aggregated embeddings. **a**. 3d renderings of representative proofread neuron and glia cell types from the mouse and human datasets. The pyramidal cell axon is cut off to fit. Not shown: 4 mouse pyramidal subtypes, 4 mouse glia subtypes, 2 human neuron subtypes. Scale bars neuronal: 100 μm; glia: 25 μm. **b**. Rendering of representative cutouts from a pyramidal dendrite (top inset) and axon (bottom inset). Different size cutouts are defined by the skeleton node aggregation radius *R*. Scale bar 100 μm. **c**. Cell type classifiers are trained on top of SegCLR embeddings after aggregation into a mean embedding over the cutout. **d**. Cell typing performance of shallow ResNet classifiers over different aggregation radiuses for the six labeled cell types in the human dataset. Zero radius corresponds to a single unaggregated embedding node. **e**. Confusion matrix for the 6-class human cell type task at 10 μm aggregation radius. **f**. Illustration of SegCLR cell type predictions over the extent of a single basket cell from the mouse test set. The orange areas are predicted correctly, while the sparse black areas show mispredictions. **g**. Cell typing performance for the mouse dataset. The 13-class task (black) uses all the ground truth labeled classes, while the 10-class task (green) combines all pyramidal cell labels into a single class. The 6-class task (blue) further reduces the neuronal labels into excitatory and inhibitory groups, comparable to the labels available on the human dataset (d). **h**. Confusion matrix for the mouse 13-class cell type task at 25 μm aggregation radius. Colored boxes indicate the group of four pyramidal cell types that were collapsed into the 10-class task, and the five excitatory and four inhibitory types collapsed into the 6-class task in (g). **Mouse neuron types**. AC: astrocyte; MGC: microglia cell; OGC: oligodendrocyte cell; OPC: oligodendrocyte precursor cell; P2-6: cortical layer 2-6 pyramidal cell; THLC: thalamocortical axon; BC: basket cell; BPC: bipolar cell; MC: Martinotti cell; NGC: neurogliaform cell. **Human neuron types**. AC: astrocyte; MGC: microglia cell; OGC: oligodendrocyte cell; OPC: oligodendrocyte precursor cell; E: excitatory neuron; I: inhibitory interneuron.

SegCLR builds on recent advances in contrastive learning of image representations^27,38^, with modifications that leverage freely-available dense automated instance segmentations of neurons and glia^12,13^. Contrastive learning methods aim to learn representations by maximizing agreement between matched (“positive”) examples in a learned latent space. SegCLR selects example pairs with respect to the segmentation: positive pairs are drawn from nearby locations (within 150 μm skeleton path length) on the same object and trained to have similar representations, while negative pairs are drawn from separate objects and trained to have dissimilar representations (Fig. 1a). We also leveraged the segmentation for input preprocessing: local 3d views of EM data, 4-5 μm on a side at 32-40 nm voxel resolution, were presented to the embedding network after being masked to feature only the segmented object at the center of the field of view (Fig. 1b, left). The network architecture was based on ResNet-18^39^, with convolutional filters extended to 3d and three bottleneck layers reducing the representation to a 64-dimensional embedding. During training, a projection head further reduced the output to 16 dimensions, on which the contrastive loss was applied ^38^ (Fig. 1b, right).

We trained SegCLR separately on two large-scale, publicly available EM connectomic datasets, one from human temporal cortex^13^ and one from mouse visual cortex^12^, that were produced via different imaging and segmentation techniques. We then inferred SegCLR embeddings with partially overlapping fields of view over all non-trivial objects (at least 1,000 voxels) within each volume. This produced a 64-dimensional embedding vector for each masked local 3d view, for a total of 3.9 billion and 4.2 billion embeddings for the human and mouse datasets respectively. SegCLR thus adds modest storage overhead relative to the full EM dataset size (human: 980 GB versus 1.4 PB at 4×4×33 nm; mouse: 1 TB versus 234 TB at 8×8×40 nm). Visualizing an illustrative subset of the resulting embeddings after dimensionality reduction via UMAP projection^40^ revealed structure across each embedding space (Fig. 1c-d). Visualizing embeddings over individual cells also revealed structure within and between them (Fig. 1e-f), suggesting the potential for embeddings to solve diverse downstream tasks.

### Cellular subcompartment classification

Embedding vectors representing local segment views throughout the EM datasets can be applied to a variety of downstream tasks, such as clustering, similarity search, or (Fig. 2a) classification. We first examined using SegCLR to distinguish cellular subcompartments such as axons, dendrites, and somas (Fig. 2b). In the human cortical dataset we also included astrocytic processes, as a distinct kind of subcompartment for which we had ground truth labeling. On a set of segmented object locations with expert labeled subcompartment identities, the respective SegCLR embeddings formed largely separable clusters in embedding space (Fig. 2c, e). A linear classifier trained to distinguish embeddings from the human cortical dataset could identify subcompartments in a held out test set with 0.988 mean F1-Score, while on the mouse dataset, axon, dendrite, and soma classification reached 0.958 mean F1-Score. The F1-Score summarizes classification accuracy and reflects both precision and recall performance via their harmonic mean.

We also tested reducing the ground truth labeling requirements, and compared the performance of subcompartment classification using SegCLR embeddings versus directly training a fully supervised subcompartment classification network^25^. The supervised network input data and network architecture (ResNet-18) were identical to the SegCLR setup, except that we replaced the SegCLR bottleneck and contrastive projection head with a classification softmax. On the 4-class subcompartment task, the embedding-based classification matches the performance of direct supervised training while requiring roughly 4,000 times less labeled training data (10-run median F1-Score, ∼400 examples total), and exceeds supervised performance when using all labeled data (Fig. 2d). We also note that for the smallest sample sizes, certain random samples performed significantly better than the average random sample (light gray points), suggesting a potential for further gains in accuracy and efficiency under more sophisticated sampling strategies^41^.

### Classification of neuron and glia subtype for large and small cell fragments

Another important application of SegCLR was for cell type classification. To assess performance, we focused on the neuron and glia types for which we had expert ground truth labels: thirteen mouse cell types and six human cell types (Fig. 3a). We also restricted the labeled set to manually proofread cells, to avoid cell type ambiguity from any residual merge errors in the reconstructions.

**Supplemental Figure 3.**
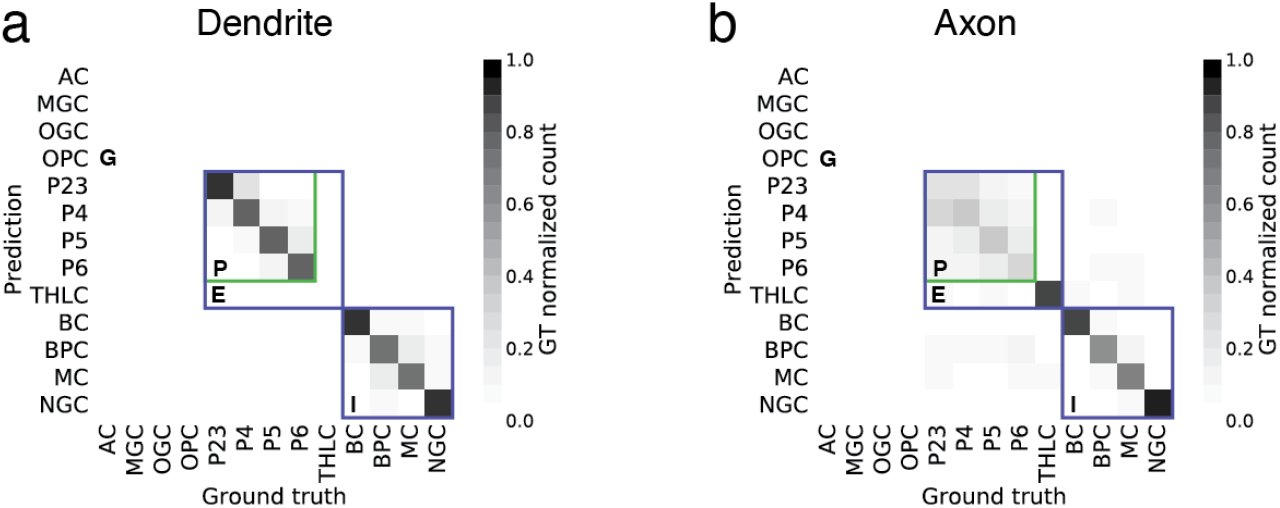
Cell type classification of large and small cell fragments via aggregated embeddings. **a**. Confusion matrix for the mouse 13-class cell type task at 25 μm aggregation radius. Colored boxes indicate the group of four pyramidal cell types that were collapsed into the 10-class task, and the five excitatory and four inhibitory types collapsed into the 6-class task in (similar to Fig. 3h). We restricted this evaluation to dendrites based on the automated compartment classification (Fig. 2). There were no ground truth examples of glia subtypes and thalamocortical cells in this analysis but the classifier could still predict those classes. **b**. As in (a), but restricted to axons. There were no ground truth examples of glia subtypes in this analysis but the classifier could still predict those classes.

While individual SegCLR embeddings representing local 3d views 4-5 μm on a side were sufficient for subcompartment classification (Fig. 2), for cell typing we found it helpful to aggregate embedding information over larger spatial extents prior to classification (Fig. 3c). Starting from a position of interest on a cell, we collected all nearby embeddings for a set distance *R* along the skeleton path in all directions. We then combined the collected set of embeddings by computing a mean embedding value over each feature dimension; this simple aggregation approach proved effective as input to the shallow two-module ResNet classifiers used for cell typing.

Ultimately we wanted to classify not only large but also small cell fragments, which are common in large-scale automated reconstructions. Therefore, an important question was how many embeddings, and over what spatial extent, need to be aggregated to achieve good cell typing. We assessed this by constructing varying sized cutouts from the ground truth cells (Fig. 3b) corresponding to different allowed aggregation distances *R*. For both mouse and human datasets, SegCLR supports high accuracy cell typing (human 6-class mean F1-Score at R=10μm 0.938; mouse 6-class mean F1-Score at R=25μm 0.947; mouse 10-class mean F1-Score at R=25μm 0.832; mouse 13-class mean F1-Score at R=25μm 0.748) at aggregation distances of only 10-25 μm (Fig. 3d-h). When using all thirteen mouse ground truth types, most residual classification errors were between pyramidal cell subtypes (Fig. 3h, Sup.Fig. 3), particularly in their axons.

### Unsupervised data exploration via SegCLR

SegCLR also proved useful for data exploration beyond supervised classification tasks. We used unsupervised UMAP projection^40^ to visualize samples of embeddings in the human and mouse datasets, and readily observed separate regions in UMAP space for glia versus neurons, and for axons, dendrites, and somas (Fig. 1c-d; Fig. 2c, e). We then focused in on smaller subregions of embedding space to reveal more fine-grained distinctions. For example, selecting embeddings representing just the dendritic arbors of pyramidal cells from cortical layer 5 in the mouse dataset revealed further detailed structure, including three main distinct UMAP clusters (Fig. 4a).

**Figure 4.**
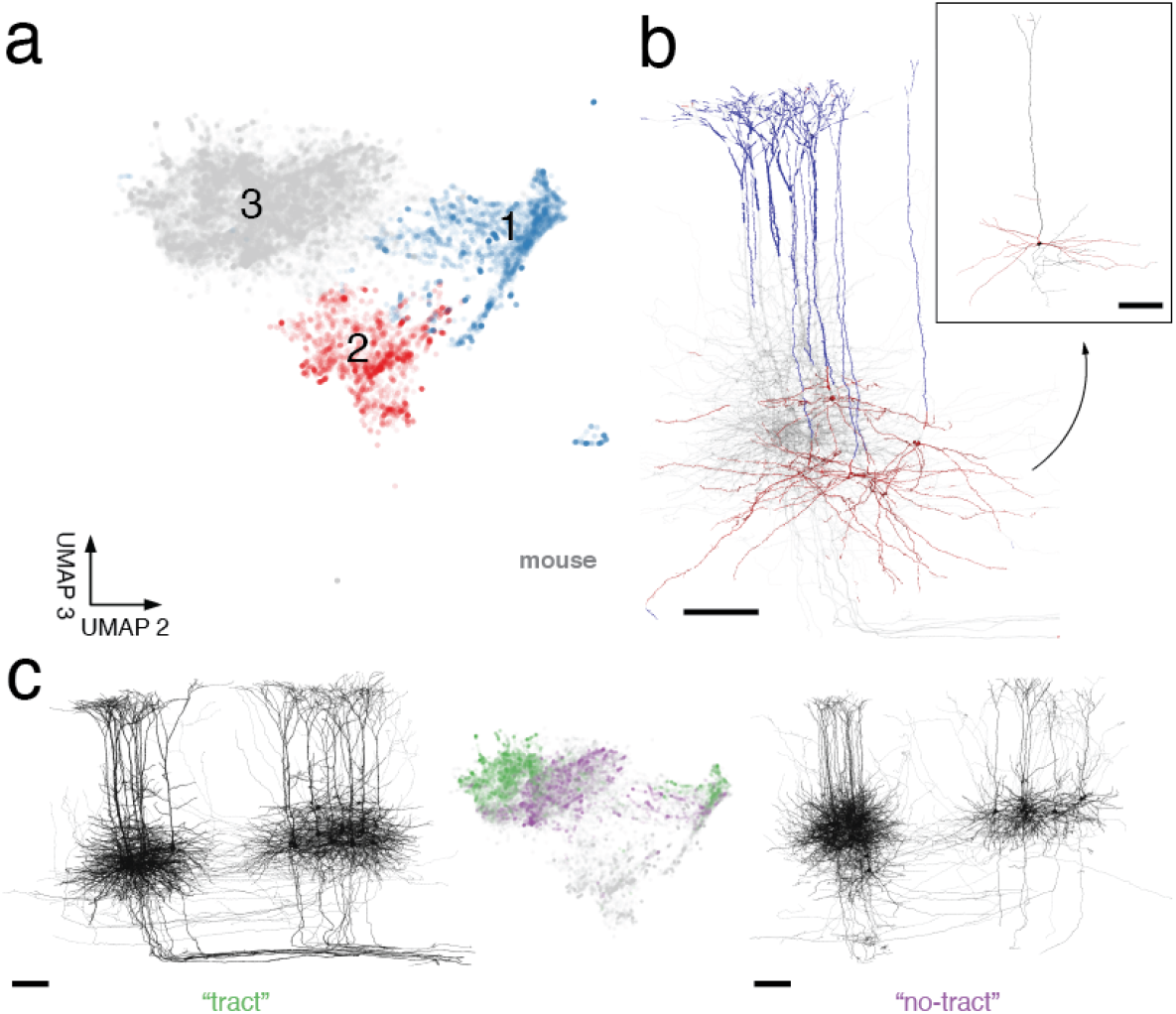
Unsupervised exploration of mouse layer-5 pyramidal dendrite embeddings. **a**. SegCLR embeddings projected to 3d UMAP space, with two selected axes displayed. Each point represents an embedding (aggregation distance 50 μm) sampled from only the dendrites of mouse layer-5 pyramidal cells. The UMAP projections separate into three main clusters. **b**. Renderings of selected cells, colored to match (a) for locations whose nearest embedding fell within cluster 1 (blue) or cluster 2 (red). Projections falling within cluster 1 are strongly associated with the apical dendrite subcompartment, while cluster 2 is strongly associated with a subset of basal dendrites corresponding to cells with a distinct “near-projecting” (NP) morphology (inset). **c**. Of the cells whose projections fall within cluster 3, a subset have distinctive axon trajectories consistent with their axons joining a major output tract (left). These “tract” cells also occupy a distinct subregion within cluster 3 (middle, green). Cells occupying the remainder of cluster 3 (middle, purple) consistently lack the axon tract morphology (right). The “tract” and “no-tract” groups are separable within both primary visual area V1 (left group of cells for both “tract” and “no-tract”) and higher visual areas (right group of cells for both “tract” and “no-tract”). Scale bars 100 μm.

Visualizing the subcellular locations whose embedding projections fell within cluster 1 revealed a strong association with the apical dendrite subcompartment (Fig. 4b), whereas locations outside cluster 1 were consistently basal dendrites or apical shafts proximal to the soma. The apical specificity of cluster 1 extended across all layer-5 pyramidal cells examined (181 total). In contrast, we found that nodes corresponding to cluster 2 were basal dendrites, and were restricted to only a small subset (13%) of cells. This subset had dendritic morphology distinct from typical pyramidal cells, featuring sparse arbors with relatively little branching, and with basal dendrites tending to descend laterally over long distances (Fig. 4b, inset).

Cluster 3 contained the basal dendrite nodes for the remaining majority (87%) of layer-5 pyramidal cells. Visualizing their entire reconstructed morphologies revealed that a subset of these cells had distinctive axon trajectories, consistent with their axons joining a major output tract (Fig. 4c, left). Interestingly, the UMAP projections for these “tract” cells occupied only one half of cluster 3 (Fig. 4c, orange points). As an inverse test, we then selected 30 cells whose projections fell in cluster 3 but were mostly excluded from the “tract” subregion (Fig. 4c, blue points). None of these 30 cells showed the axon tract morphology (Fig. 4c, right). Thus, SegCLR embeddings of basal dendrites largely separate two pyramidal subgroups that independently separate based on their axon trajectories.

Comparison of our grouping of layer-5 pyramidal cells to previous work suggests that, based on its distinct morphology and relative rarity, our cluster 2 group (Fig. 4b, inset) corresponds to the type described in the literature as near-projecting (NP) or cortico-cortical non-striatal (CC-NS)^42–44^ cells. Consistent with NP basal dendrites being sparse and more morphologically similar to apical dendrites than in typical pyramidal cells, the NP basal cluster 2 was also closer to the apical cluster 1 in SegCLR UMAP space.

Our cluster 3 “tract” group (Fig. 4d) likely corresponds to the type described in the literature as extra-telencephalic (ET), cortico-subcortical (CS), thick-tufted (TT) or pyramidal tract (PT), which provide the primary cortical output onto subcortical brain areas^42–44^. The cluster 3 “no-tract” cells (Fig. 4f) then likely correspond to the intra-telencephalic (IT) or cortico-cortical (CC) type. While the NP, ET, and IT subtypes have previously been distinguished by gene expression, developmental history, electrophysiology, morphology, or their distal projection targets, we add evidence that they can be distinguished by their dendritic morphology and proximal axonal trajectory. It is also notable that the separate ET and IT groups within cluster 3 are maintained across both the primary visual area (V1) and higher visual areas (HVA) included in our dataset (Fig. 4d-f).

### Out-of-distribution input detection via Gaussian processes

A remaining issue with applications such as cell typing on large-scale datasets is how to gracefully handle image content that falls far outside the distribution of labeled examples. These “out-of-distribution” (OOD) input examples could include locations containing imaging artifacts or segmentation merge errors, or they could represent genuine biological structures that were simply absent in the training set. One example of the latter case is the common analysis situation in which only a few cell types have confident ground truth labels, but one wishes to classify these types while avoiding spurious classifications of a potentially large majority of surrounding segments belonging to diverse unknown types.

We addressed OOD inputs via Spectral-normalized Neural Gaussian Processes^34^ (SNGP), which add a prediction uncertainty to the model output (Fig. 5a), and calibrate that uncertainty to reflect the distance between the test data and the training distribution. This allows OOD inputs to be detected and rejected, rather than spuriously classified, while requiring no extra labeling effort. To evaluate SNGP, we constructed a human cortical cell type dataset in which only the glial labeled types were used to train classifiers, while both glial and neuronal types were presented for testing (Fig. 5b). The neuronal types, making up 50% of the constructed test set, thus served as an OOD pool.

**Figure 5.**
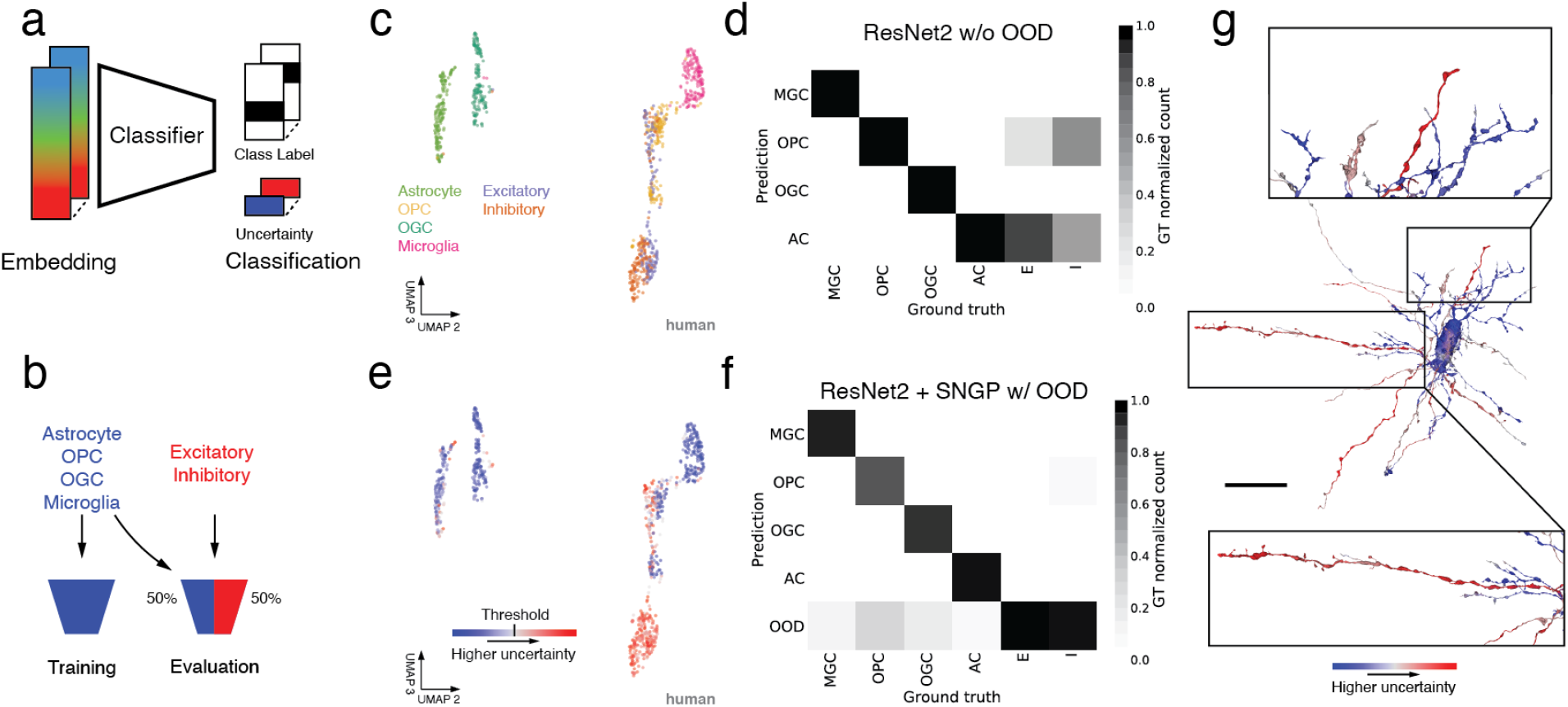
Out-of-distribution input detection via Gaussian processes. **a**. We handled OOD inputs by computing prediction uncertainties alongside class labels, and calibrated the uncertainties to reflect the distance between the test example and the training distribution. **b**. To evaluate OOD detection, we trained classifiers on glial cell type labels, and then evaluated the classifiers on a 50/50 split between glial and OOD neuronal cell types. **c**. UMAP projection of locally aggregated embeddings (radius 10 μm) from the human cortical dataset, colored by ground truth labeled cell type. **d**. Confusion matrix for a ResNet-2 classifier trained on only the four glia types, with OOD neuronal examples mixed in at test time. While in-distribution glial classification is almost flawless, all neuronal examples (half the constructed test set) receive spurious classifications. **e**. As in (c) with the UMAP projected embeddings now colored by their SNGP uncertainty. The colormap transitions from shades of blue to shades of red at the threshold level used to reject OOD in our experiment. **f**. Confusion matrix for the SNGP-ResNet-2, assembled from five-fold cross-validations. Examples that exceed the uncertainty threshold are now treated as their own OOD predicted class. The strong in-distribution glia performance of (d) is largely retained, while neuronal fragments are effectively filtered out. **g**. Spatial distribution of local uncertainty over an unproofread segment that suffers from reconstruction merge errors between the central OPC glia and several neuronal fragments. The uncertainty signal distinguishes the merged neurites (red: high uncertainty / OOD) from the glia cell (blue: low uncertainty) with a spatial resolution of roughly the embedding aggregation distance. Scale bar 25 μm. See Fig. 3 for cell type abbreviations.

We first trained a small conventional network (i.e., lacking SNGP capabilities) on the glial classification task. Specifically, a shallow two-module ResNet classifier (“ResNet-2”) was trained on locally aggregated embeddings (radius 10 μm) from only the glial labeled cells. This network performed with high accuracy on the in-distribution glial half of the test set, but also (by construction) spuriously classified all OOD neuronal examples (Fig. 5d). The goal of OOD detection is to retain the strong in-distribution performance while selectively filtering out OOD inputs. To achieve this we converted the ResNet-2 to a SNGP by making drop-in substitutions to the trained network’s hidden and output layers^34^. These changes equip the classifier with an uncertainty output that estimates in part the degree to which each example is OOD with respect to the training distribution (Fig. 5e). The SNGP uncertainty can then be thresholded at a task-appropriate level to determine how aggressively to reject OOD inputs.

The SNGP-ResNet-2 retains strong in-distribution glia classification performance while effectively filtering out OOD neuronal examples (Fig. 5f). With uncertainty thresholded at a level that optimizes F1 on a validation set and the resulting OOD examples treated as a separate class, overall mean F1-Score reaches 0.888. Note that the network layer substitutions for SNGP apply only to the small classifier network, with no modifications required to the underlying SegCLR embeddings. Furthermore, the neuronal ground truth labels used here were only needed to validate the results, while to train and deploy a classifier with SNGP OOD detection requires no extra ground truth labeling beyond the in-distribution set. These results show that SegCLR can be effectively deployed in common settings where only a small selection of expert labels may be available for application to a dataset containing a large proportion of OOD objects. We also note that when used with SegCLR, OOD classification may prove simpler to implement than in non-embedding settings (see Methods).

Finally, we also evaluated the spatial distribution of local uncertainty over larger segments. This is particularly relevant for unproofread segments that contain reconstruction merge errors between a labeled and an OOD type. For example, the uncertainty of our SNGP classifier can distinguish neuronal fragments erroneously merged onto a central glia (Fig. 5g). Because the embeddings are aggregated with a local 10 μm radius, the distinction between neuronal and glial branches of the segment can be resolved to within roughly 20 μm. Automated merge error correction based on these branch distinctions^13,25^, potentially combined with direct detection of merge-specific features from embeddings at the merge point^45^, would be an attractive application for future investigation.

### SegCLR cell typing of pre- and post-synaptic partners for large and small fragments

In brain circuit analysis, a common goal is to identify the cell types of the thousands of synaptic partners upstream or downstream of a particular cell or circuit of interest^46–48^. Due to the incompleteness of current automated reconstructions, contemporary connectomics-based circuit analysis efforts typically require significant manual tracing to extend each partner neurite back toward its soma in order to provide enough morphological structure to enable an expert to propose a cell type identity. Furthermore, in many datasets partner neurites may terminate at a volume boundary before their cell type is resolvable. However, as demonstrated here (Fig. 3), SegCLR is able to classify cell types of even relatively small cell fragments with high accuracy, enabling large-scale analysis of synaptic partners without manual tracing.

We leveraged this new capability to perform detailed analysis of synaptic partners across multiple cell types in mouse cortex (Fig. 6). For each synapse annotated in this dataset^12^ we gathered the predicted cell type annotation of the closest embedding nodes pre- and postsynaptically. We filtered out 9.2% of presynaptic and 7.8% of postsynaptic fragments that were too short (R_max_ < 2.5 μm, see methods) and ignored a further 8.8% and 2.5% respectively because of high uncertainty scores.

First, we analyzed the distribution of presynaptic cell types for a representative sample of 919,500 synapses at varying cortical depths (Fig. 6a). Our method classified 72.3% of presynapses and 76.3% postsynapses as excitatory (ignoring those marked as uncertain). We then focused on a core set of proofread cells (e.g. Fig. 6b), for which we collected all their input and output synapses, along with the corresponding (unproofread) presynaptic and postsynaptic partner fragments. For each proofread cell, we then analyzed the relative proportions of received inputs by cell type (Fig. 6c-e) as well as the proportion of downstream targets (Fig. 6h-l).

**Figure 6.**
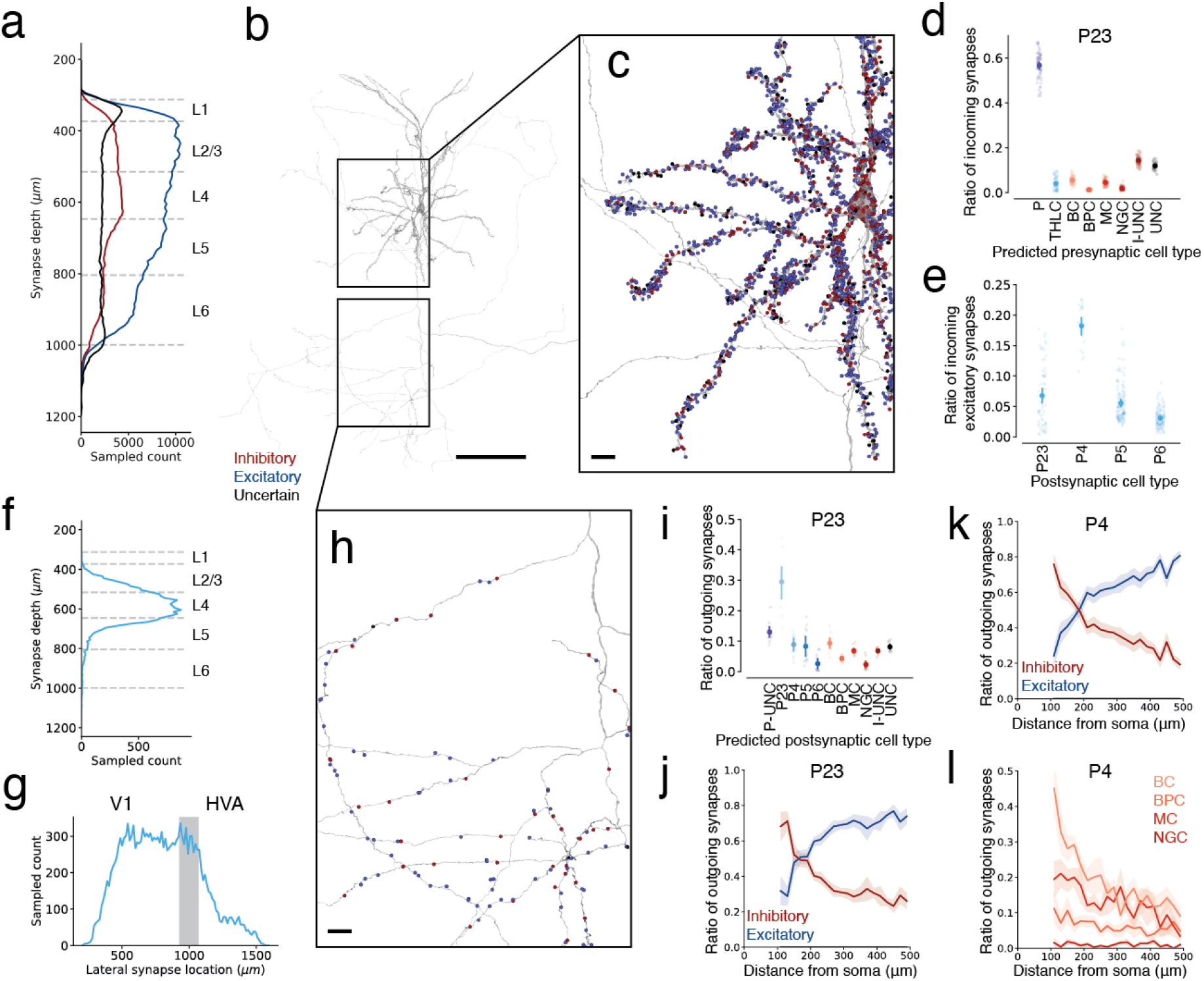
Quantitative analysis of pre- and post-synaptic partner cell type frequencies. **a**. The distribution of synapses by predicted inhibitory versus excitatory presynaptic axonal cell type, over the depth of the mouse visual cortex. **b**. Representative layer 2/3 pyramidal cell. **c**. Pyramidal cell input synapse locations, annotated (dots) and colored by predicted inhibitory versus excitatory presynaptic partner cell type. **d**. Distribution of upstream presynaptic partners for layer-2/3 pyramidal cells in V1 with proofread dendrites (N=69). Each cell is represented by a set of points describing its ratios of input cell types. The mean and standard deviation of the ratios are also shown (darker point and line). UNC refers to synapses where no classification could be made. I-UNC refers to synapses where only a coarse classification call (excitatory versus inhibitory) could be made. **e**. The ratio of thalamocortical versus pyramidal innervation onto pyramidal targets in different cortical layers shows major thalamic input to cortical layer 4 (N: P23=69, P4=23, P5=136, P6=127). **f**. The thalamocortical synapse counts over the cortical depth, showing concentration in layer 4. **g**. The thalamocortical synapse counts along the lateral axis through the dataset drops at the V1-HVA boundary. The shaded area shows the approximate projection of the V1-HVA border onto the lateral axis. **h**. As in (c), for downstream postsynaptic partners **i**. As in (d), distribution of downstream postsynaptic partners for layer-2/3 pyramidal cells in V1 with proofread axons (N=12). P-UNC refers to synapses where SegCLR was not able to classify a pyramidal subtype with sufficient certainty. **j.k**. Inhibitory versus excitatory balance of downstream postsynaptic partners with increasing distance along P23 axons (N=12) (j) and P4 axons (N=19) (k). Mean and standard error of the mean within each distance bucket are shown. **l**. As in (k), broken up into individual inhibitory subtypes (N=19).

**Supplemental Figure 6.**
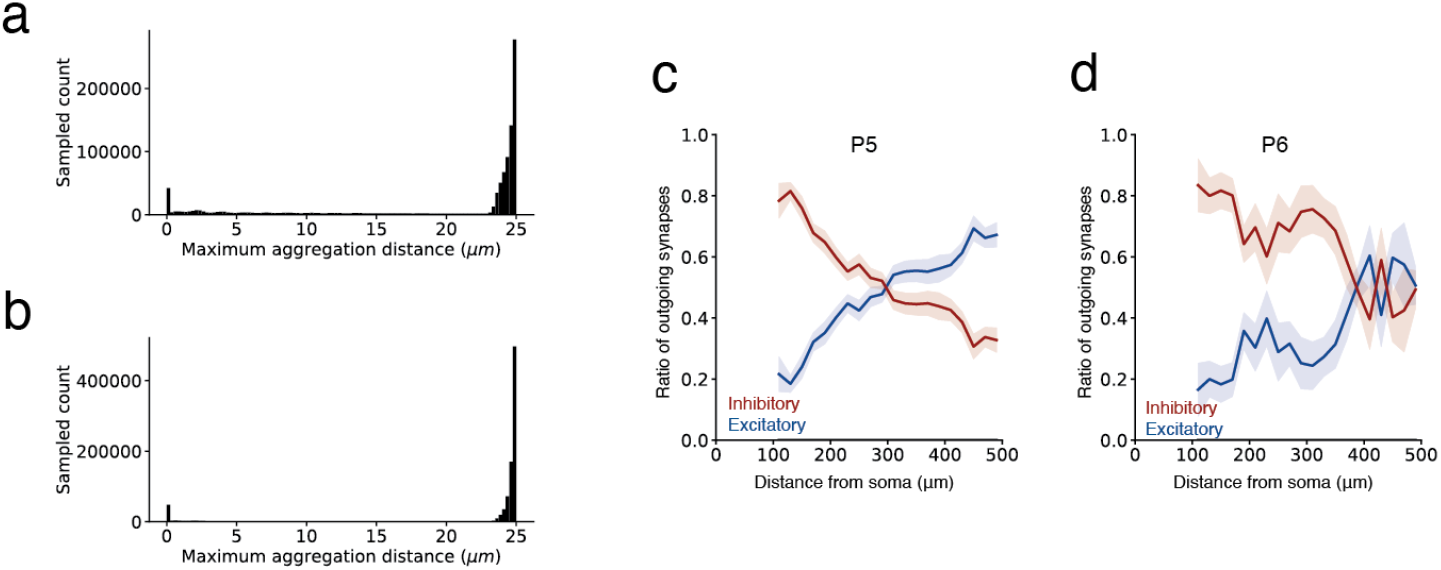
Quantitative analysis of pre- and post-synaptic partner cell type frequencies. **a**. Distribution of the distance between the farthest embedding node and the queued embedding node (R_agg_=25 μm) for presynaptic segments. In most cases, the distance to the farthest embedding node is close to the maximal permitted distance, in a few cases the presynaptic segment was small with only a few embedding nodes. **b**. As in (a) for postsynaptic segments. **c.d**. Inhibitory versus excitatory balance of downstream postsynaptic partners with increasing distance along P5 axons (N=34) (c) and P6 axons (N=6) (d).

For upstream (presynaptic) partners, we found significant differences in the proportion of intracortical (from pyramidal cells) versus subcortical (from putative thalamocortical axons) excitatory inputs (Fig. 6e-g). In layer 4, the canonical cortical input layer, pyramidal cells showed significantly more subcortical innervation, with the putative thalamocortical synapses contributing 18% of their excitatory inputs (Fig. 6e-f). This percentage agrees with estimates based on immunohistochemistry in mouse V1^49^ and other mouse cortical regions^50^. When plotting the prevalence of layer 4 thalamocortical synapses along an axis from V1 to HVA (higher visual areas), however, we observed a drop in thalamocortical innervation that coincided with the region boundary (Fig. 6g). Thalamocortical projections to HVA had been identified previously^51–53^ but had not been quantified, demonstrating how the proposed computational approach can give new quantitative insights into cortical cell type connectivity.

For downstream partners (Fig. 6h-l), we analyzed the relative proportion of output synapses onto excitatory versus inhibitory partners as a function of distance along the presynaptic axon (Fig. 6k-l, Sup.Fig. 6 c,d). Across all cortical layers, the distribution of output synapses along pyramidal cell axons was biased towards more inhibitory postsynaptic partners in the proximal regions^54,55^, with excitatory downstream partners growing more common with increasing distance from the soma^56^. Our analysis is thus broadly consistent with previous reports for layer 2/3 visual^54,55^ and entorhinal cortex^56^, but further showed details of axonal sorting in layer 4 and revealed that changes in inhibitory targeting over axonal distance are driven primarily by declines in targeting of Basket and Martinotti cells.

Using SegCLR cell type predictions for pre- and post-synaptic fragments, we thus derived results consistent with prior reports that relied on laborious manual annotation, while extending their scale in terms of number of synapses, cell types, and brain areas analyzed.

## Discussion

We have introduced SegCLR, a self-supervised method for training rich representations of local cellular morphology and ultrastructure, and demonstrated its utility for biological annotation in human and mouse cortical volumes. Beyond the requirement for an accompanying instance segmentation, the current SegCLR formulation has some limitations. First, the 32-40 nm voxel resolution of input views impedes capture of finer EM ultrastructural features, such as vesicle subtypes or ciliary microtubule structure. Training SegCLR on higher resolution inputs, or with multiscale capabilities, is worth detailed exploration.

Another limitation is that explicit input masking excludes EM context outside the current segment, while in some cases retaining surrounding context could be useful, e.g. for myelin sheaths, synaptic clefts, and synaptic partners. We therefore tested a version of SegCLR that receives the unmasked EM block and the segmentation mask as two separate input channels, rather than a single explicitly masked EM input. This variant performed similarly on the subcompartment classification task, but appeared more sensitive to subtle nonlinear photometric differences across the extent of the dataset. Given SegCLR’s ability to reliably classify cell types that human experts find challenging to distinguish, including for small cellular fragments, exploration of attribution methods^57^ to reveal which SegCLR features are decisive could inform future automated methods as well as human understanding.

SegCLR currently also focuses only on a single segment, whereas it could be useful to additionally learn representations of dual- or multi-cellular complexes. Training SegCLR on multi-segment inputs is conceptually straightforward, but running the network on all possible segment pairs or complexes within a large-scale dataset would be prohibitive. A strategy for limiting the set of targeted complexes, e.g. only using predefined synaptic partners, would be needed.

Finally, the local 4-5 μm input field of view for each embedding could be considered a limitation; many tasks, including cell typing, are expected to benefit from larger contexts^28,30,31^. In the current work, we demonstrated that a simple mean embedding aggregation strategy is sufficient for reasoning over larger contexts. However, more sophisticated aggregation methods^28,58,59^ could still prove useful for generating representations of larger contexts. There are also opportunities to extend representations beyond single cells. For example, neuronal embeddings could be extended with additional dimensions aggregated from pre- and post-synaptic partners, to create connectivity-enhanced cell type signatures or to form motif representations.

We expect SegCLR to be widely applicable across the growing breadth of volumetric EM datasets, and speculate that it could also apply in other settings where available segmentations can serve as a guide to representation learning^60–63^. While the computational demands for training and deploying embeddings are considerable, this cost is incurred only once, rather than repeatedly for each downstream analysis. We will make Python code available for input preprocessing, network training, and evaluation, and will release pretrained TensorFlow network weights.

By providing rich and tractable representations of EM data, SegCLR greatly simplifies and democratizes downstream research and analysis. For example, the previous state of the art in subcompartment classification required millions of training examples assembled from thousands of manually validated segments, thousands of GPU hours to train a deep network classifier, and hundreds of thousands of CPU hours to evaluate over a large-scale dataset^13,25^. With SegCLR embeddings in hand, this benchmark is outperformed by a simple linear classifier, trained in minutes on a single CPU, with a few hundred manually labeled examples (Fig. 2).

Arguably the most powerful application of SegCLR demonstrated here is the ability to classify neuronal and glial subtypes even from small fragments of cells (Fig. 3). This capability has important ramifications, particularly for datasets where reconstructed cells are incomplete, or where only a portion of the brain tissue was imaged. Being able to identify connectivity patterns between specific cell types is fundamental to interpreting large-scale connectomic reconstructions. We showed how SegCLR enables cell typing of synaptic partners across large areas of mouse visual cortex, yielding new insights into e.g. the regional specificity of thalamocortical inputs and the balance of outputs onto inhibitory versus excitatory circuits over the length of pyramidal cell axons (Fig. 6).

We release the full SegCLR embedding datasets for the <u>human</u> and <u>mouse</u> cortical volumes to the community, to enhance exploration and understanding of these rich and complex resources.

## Methods

### Datasets

We used two large-scale serial-section EM connectomic datasets for SegCLR experiments: one from the human temporal cortex, imaged via SEM^13^; and one from the mouse visual cortex, imaged via TEM^12^. Both datasets are freely available and provide an aligned and registered EM volume with an accompanying automated dense instance segmentation. For SegCLR experiments we downsampled the human and mouse data to 32×32×33 nm and 32×32×40 nm nominal resolution respectively. The human EM volume was also CLAHE normalized^64^. We skeletonized both segmentation volumes as previously described^13,35^.

For subcompartment classification (Fig. 2), ground truth human data was collected on an earlier pre-release version of the dataset, as well as two smaller cutouts^13^. All three regions are contained within the publicly released dataset, although they differ slightly in their alignment and photometric normalization. For evaluation of local cell type classification (Fig. 4), it was important to have cells with minimal reconstruction merge errors in the ground truth labeled set. We therefore proofread some cells in the human dataset to exclude regions close to observed merge errors from the embeddings cell type analysis (Figs. 3-4). On the mouse dataset, we restricted analysis to the set of ground truth labeled cells that were already expert proofread prior to release^12^.

### Training SegCLR embedding networks

SegCLR was inspired by SimCLR^27,38^. The embedding network was a ResNet-18 architecture^39^ implemented in TensorFlow, with convolutions extended to 3d and 3 bottleneck layers prior to a 64-dimensional embedding output. During training, we added 3 projection layers prior to a normalized temperature-scaled cross entropy (“NT-Xent”) loss^38^ with temperature 0.1. The total number of model parameters was 33,737,824.

We also found that downstream task performance was enhanced by adding a decorrelation term to the loss, defined as:

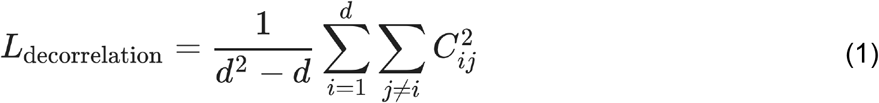

where *d* is the embedding dimensionality and *C* is the correlation matrix between embeddings over the batch. We trained SegCLR networks on 8×8 v2 Cloud TPUs for up to 350,000 steps with a full batch size of 512 pairs (1024 individual examples) and learning rate decay schedule starting at 0.2. Separate networks were trained for the human and mouse datasets. Training took about one week and proceeded at a speed of 0.5-0.6 steps/s.

The input to the network was a local 3d cutout of EM data 129 voxels on a side, nominally corresponding to 4128×4128×4257 nm in the human EM dataset, and 4128×4128×5160 nm in the mouse dataset. We then masked the EM data by the segmentation for the object at the center of the field of view, so that possible confounds in the surrounding EM context would be excluded.

We also leveraged the segmentation and corresponding skeletonization to generate example pairs for contrastive training. For an arbitrary segment, we picked a 3d view to be centered on an arbitrary skeleton node, and then picked a positive pair location centered on a second node within 150 μm path length away on the same skeleton. Positive pairs were preprocessed before training for higher performance. As there are more possible pairs for larger distances, we sorted these positive pairs into four distance buckets from which we drew uniformly. The bucket boundaries were (0, 2500, 10000, 30000, 150000) nanometers.

As in SimCLR^27^, we used the 510 examples from the rest of the batch, which were drawn from 255 other segments throughout the volume, as negative pairs. We also applied random reflections, and photometric augmentations^21^ to the inputs, to prevent the network from solving the contrastive task via trivial cues such as the orientation of processes, or the local voxel statistics.

### SegCLR inference

We inferred SegCLR embeddings over the full extent of the human and mouse datasets. After removing trivial segments, we extracted local 3d views centered on skeleton nodes for the remaining segments, with approximately 1500 nm path length spacing. The set of views for each segment therefore had substantial overlap of about 65-70% with typical nearest neighbor views. We then ran SegCLR on all selected views (3.9 billion and 4.2 billion for the human and mouse datasets respectively) via an Apache Beam Python pipeline running on a large CPU cluster, and stored the resulting embedding vectors keyed by segment ID and spatial XYZ coordinates. Inference proceeded at about 1 example/s/CPU with a batch size of 1. For visualization, we ran UMAP dimensionality reduction^40^ to 2-4 dimensions on representative samplings or on subsets of interest from among the embeddings. When sampling from a large population of embeddings of local cutouts, we sampled such that all classes were represented with a significant number of examples.

### Subcompartment classification

We trained linear classifiers to identify subcompartment types from embedding inputs based on expert ground truth labels on each dataset (Table 1). For comparison, we also trained a fully-supervised subcompartment classifier directly on voxel inputs using an identical 3d ResNet-18 architecture and input configuration (photometric augmentation was omitted and random 3d rotations were added), and replaced the bottleneck, projection heads, and contrastive loss with a classification softmax and cross-entropy loss. The supervised network was trained on 8 GPUs with a total batch size of 64 via stochastic gradient descent with learning rate 0.001 for 1.5 M steps. During full supervised training, examples were rebalanced class-wise by upsampling all classes to match the most numerous class. However, in SegCLR experiments (Fig. 2) we showed that performance using embeddings was robust to substantial reductions in training data via random sampling of examples. During subsampling we ensured that every class was represented with at least 10% of the examples. We repeated each sampling round 20 times for sample sizes ≤ 5000.

**Table 1.** Cellular subcompartment ground truth label sets. The number of distinct segments and total number of training examples of each type available for subcompartment classification (Fig. 2). For fully supervised training on the human dataset, the full set of training examples was used, while for linear classifiers trained on SegCLR embeddings, both full and reduced subsets of the training set were evaluated (Fig. 2d).

| dataset | subcompartment | #training segments | total #training examples |
| --- | --- | --- | --- |
| human | axon | 1,256 | 262,395 |
|  | dendrite | 933 | 1,151,175 |
|  | soma | 993 | 40,649 |
|  | astrocytic process | 212 | 1,392,702 |
| mouse | axon | 121 | 4,653,326 |
|  | dendrite |  | 2,343,169 |
|  | soma |  | 13,830 |

### Cell type classification

We tested cell type classification using a set of expert ground truth labeled neurons and glia from both human and mouse (Table 2). These ground truth cells are generally large and contain somas within the volume, but they are not necessarily completely reconstructed. We manually proofread human ground truth cells for merge errors by marking bad agglomeration edges in the agglomeration graph prior to evaluation. In the mouse dataset, we restricted analysis to proofread neurons included in the public v343 release. We trained shallow 2-module ResNets to predict cell subtypes (Fig. 3, Sup Fig. 3).

**Table 2.** Cell type ground truth label sets. The number of distinct segments and total number of embeddings available for cell type classification (Figs. 3-6). Note that only a fraction of the embedding nodes were used during training.

| dataset | cell supertype | cell subtype and abbreviation | #segments | #embeddings total |
| --- | --- | --- | --- | --- |
| human | neuron | Excitatory (E) | 159 | 863,644 |
|  |  | Inhibitory (I) | 52 | 91,283 |
|  | glia | Microglia cell (MGC) | 36 | 29,536 |
|  |  | Oligodendrocyte precursor cell (OPC) | 19 | 23,614 |
|  |  | Oligodendrocyte glia cell (OGC) | 17 | 4,449 |
|  |  | Astrocyte (AC) | 45 | 1,394,056 |
| mouse | neuron | Layer 2/3 Pyramidal cell (P23) | 95 | 1,281,244 |
|  |  | Layer 4 Pyramidal cell (P4) | 23 | 254,156 |
|  |  | Layer 5 Pyramidal cell (P5) | 36 | 633,328 |
|  |  | Layer 6 Pyramidal cell (P6) | 8 | 67,504 |
|  |  | Basket cell (BC) | 76 | 1,347,142 |
|  |  | Bipolar cell (BPC) | 33 | 141,179 |
|  |  | Martinotti cell (MC) | 30 | 460,301 |
|  |  | Neurogliaform cell (NGC) | 17 | 126,407 |
|  |  | Thalamocortical axon (THLC) | 11 | 108,639 |
|  | glia | Microglia cell (MGC) | 9 | 22,819 |
|  |  | Oligodendrocyte precursor cell (OPC) | 6 | 16,404 |
|  |  | Oligodendrocyte glia cell (OGC) | 28 | 20,142 |
|  |  | Astrocyte (AC) | 6 | 269,367 |

For cell typing, we aggregated local embeddings by collecting all the embedding nodes within a 0-50 μm radius window along the cell’s skeleton path length. A simple aggregation method of taking the mean embedding value over each feature dimension performed well when using shallow ResNets for the downstream task (Figs. 3, 5, 6) or for unsupervised data exploration (Fig. 4).

We evaluated classification performances by randomly selecting 75% of cells for each cell type for training and 25% for testing. We repeated this sampling 10 times for each aggregation distance and computed the mean across all runs. Within each run we computed the F1-Score of each class individually and then combined them for an overall mean F1-Score. In each run we sampled 30,000 (mouse) or 100,000 (human) embeddings from the cells in the training set, equally distributed across all cell types. During testing, imbalances between classes were balanced by repeating examples from minority classes. The confusion matrices in Figure 3 were generated by concatenating the test sets of all runs of one aggregation distance first. We then sampled 100,000 test examples equally distributed across all cell types. For the mouse dataset, we then repeated this procedure but restricted the sampling to nodes that were automatically labeled as dendrite or axon (Sup. Fig. 3).

To obtain results for 10 and 6 classes (Fig. 3g) we added up the probabilities of the combined classes. For instance, we added up the probabilities for all pyramidal subtypes to a single pyramidal class for the 10-class evaluation. We then followed the same procedure as for the 13-class evaluation.

### Cell-type ground truth

The different neuronal types in the mouse dataset were classified based on the morphological and synaptic criteria as described^10^. Pyramidal cells were identified by the presence of a spiny apical dendrite radiating toward the pia, spiny basal dendrites and an axon that formed asymmetric synapses. Putative thalamic axons also formed asymmetric synapses and though their soma was located outside the reconstructed volume, their gross morphology resembles previously described thalamic arbors^65^ and their fine morphology at the ultrastructure levels^50^. Neuronal cells were classified as inhibitory interneurons if their axon formed symmetric synapses. Inhibitory interneurons were further assigned subtypes using their synaptic connectivity and the morphology of axons and dendrites^66–68^. Basket cells were identified by having a larger number of primary dendrites, and at least 12% of their postsynaptic targets were pyramidal cell somata. Martinotti cells were identified by its apical axon that projected to cortical layer 1 and targeted mostly distal dendritic shafts and spines of excitatory cells; consistent with^66^. Martinotti cells also were characterized by having a multipolar dendritic arbor that was usually spiny. Bipolar cells usually had two to three primary dendrites. The dendrites were usually spiny and showed a vertical bias. Neurogliaform cells were often (but not exclusively) in cortical layer 1. The axons of neurogliaform cells have usually lower density of synapses; consistent with^66^. Neurogliaform cells also had a large number of primary dendrites and have a different pattern of synaptic inputs when compared with other inhibitory cell types.

### Unsupervised exploration of mouse visual cortex layer-5 pyramidal cells

For the detailed exploration of mouse layer-5 pyramidal cells (Fig. 4) we selected only the embedding nodes that were subcompartment classified as dendrites (Fig. 2) from a set of cells that were dendrite proofread and labeled as layer-5 pyramidal cells by human experts (N=181). We then randomly sampled up to 1051 embeddings per cell for a total of 146,607 and ran unsupervised UMAP^40^ to project the embeddings to three dimensions using Manhattan distance, 100 neighbors, 500 epochs, and learning rate 0.5.

To semi-automatically collect the projections corresponding to the visualized 3d UMAP clusters (Fig. 2a), we ran *k*-means++ with 25 groups and manually identified which *k*-means groups corresponded to UMAP clusters 1, 2, and 3. We found that running *k*-means on the raw embedding space suffered from the influence of spurious dimensions, such as dimensions that primarily picked up subtle differences in the image statistics of groups of z-sections that were captured on different microscopes^12^. On the other hand, we found that *k*-means directly on 3d UMAP struggled to capture the visual clusters due to their somewhat irregular shapes. Therefore, we found *k*-means on a rerun 5d UMAP, with the same 146,607 input embeddings and with other parameters unchanged, to be most effective.

Selecting cells with greater than 40% of their dendrite projections in cluster 2 (N=24) isolated the near-projecting pyramidal subtype (Fig. 2b, inset). To isolate the “tract” (putative extra-telencephalic) subset, we rendered all the cells with significant cluster 3 occupancy, equivalent to those with less than 25% of their projections in cluster 2 (N=157), and then manually selected the subset with distinct output tract axon trajectories (N=19; Fig. 2c, left). This group likely misses some extra-telencephalic cells in our set due to incomplete axon reconstructions, but it clearly identifies a “tract” subregion of cluster 3 (Fig. 2c, middle) and corresponding “tract” *k-*means groups. Finally, to isolate the “no-tract” (putative intra-telencephalic) subset, we scored all cluster 3 cells based on relative occupancy of “tract” *k*-means clusters, and selected a subset (N=30) that was least “tract” weighted. Visualizing these cells showed that none of them had the “tract” axon morphology (Fig. 2c, right), providing an inverse confirmation of the clustering.

### Out-of-distribution input detection

We detected OOD (OOD) inputs via Spectral-normalized Neural Gaussian Processes^34^ (SNGPs). As a baseline we trained a shallow ResNet-2 (2 ResNet modules) to classify glial cell types of 10 μm radius fragments. We then modified the ResNet-2 by spectrally normalizing its hidden layers and replacing the output layer with a Gaussian process, using the SNGP package in TensorFlow. We used the BERT hyperparameter setting from Liu et al.^34^ for our analysis.

We evaluated performance on a test set constructed of a 50/50 split between glia and OOD neuronal examples. First, we computed prediction uncertainty estimates for the test networks using the Dempster-Shafer metric^34^:

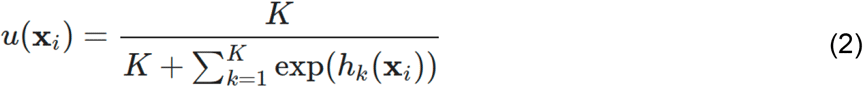

where *K* is the number of classes (in our case: 4) and *h*_*k*_ are the classification logits (network outputs prior to probability normalization). As suggested in the SNGP tutorial, we replaced the Monte Carlo estimation of the output logits with the mean-field method ^69^:

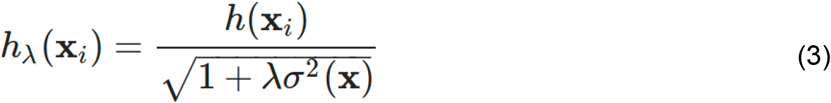

using λ = 3/π^2^ and variances estimated with the GP module.

For the evaluation, we repeated the 5-fold cross-validation as outlined for the cell type fragment classification. We sampled 100,000 unique examples and upsampled them such that each glia class accounted for 12.5% and each OOD class (excitatory, inhibitory) accounted for 25%. During each fold we trained a new classifier on a subset of the glia examples and then predicted the examples in the hold-out test set as well as the set of neuronal fragments which we reused for all 5 folds. For each fold we found the uncertainty threshold that maximized the F1-Score of the in-distribution vs out-of-distribution task. For this, we set aside half the examples from the test-set which were then not used for calculating the scores. For each fold, we replaced the original class prediction (1 of 4 glia classes) with the OOD class when the uncertainty for an example exceeded this threshold. Finally, we calculated F1-Scores for each of the 5 classes after scaling them such that OOD examples accounted for 50% and averaged them for a final F1-Score per fold. We reported the mean F1-Score across the five folds.

### Automated analysis of synaptic partners

We applied the best performing 25μm ResNet2+SNGP model from our 13-class cell type classification (Fig. 3) as well the same model restricted to 3-classes to all embeddings in the dataset. While classifications from the 13-class model can be aggregated to a 3-class classification, we were interested in the uncertainties produced by the 3-class model to filter out uncertain examples. Here, we λ = 2 to compute adjusted logits (3).

First, we labeled nodes as uncertain where the furthest aggregation distance was below 2.5μm indicating that the segment was very small (Sup. Fig. 6a,b) oftentimes only containing a single embedding node. We note that this affects more nodes close to the dataset boundaries where data quality is lower (see Fig. 6a). Next, we labeled nodes as uncertain where the predicted uncertainty (2) was above 0.45. For the remaining nodes, we assigned the label with the highest classifier probability. Some nodes were predicted to belong to the glia class; for the analysis in Fig. 6 we assigned these to the uncertain category.

Next, we attempted to assign sub-type labels to nodes classified as inhibitory or excitatory. For inhibitory nodes, we assigned the subtype with the highest predicted probability when the predicted uncertainty was below 0.05. We assigned a generic I-UNC label for all remaining inhibitory nodes. For excitatory nodes, we assigned subtypes when the predicted uncertainty was below 0.05. Here, we assigned a thalamocortical label when the predicted thalamocortical probability exceeded the summed probability across all pyramidal cell types and the pyramidal subtype with the highest predicted probability otherwise. We assigned a generic P-UNC for all remaining excitatory nodes.

We assigned cell type classifications to synapses by finding the closest embedding nodes in euclidean space pre- and postsynaptically and using their cell type labels.

To analyze cell type distributions across the dataset, we randomly sampled 919,500 synapses from the entire dataset. We corrected the native coordinates to orient the dataset vertically between white matter and pia using the standard_transform package (https://github.com/ceesem/standard_transform).

### Axonal sorting

We skeletonized all cells used in the analysis to obtain distances between synapses and the respective soma along the axon. We assigned a distance to each synapse and binned synapses with a width of 20 μm for which we computed the ratio of synapses predicted as excitatory and inhibitory (Fig. 6j,k) or their respective subtypes while ignoring synapses predicted as uncertain. For each bin we computed the mean and standard error of the mean across all cells included in the analysis.

### Embedding datasets

We make the embeddings for the mouse and human datasets available as sharded csv files. mouse: *gs://iarpa_microns/minnie/minnie65/embeddings_m343/segclr_csvzips/README* human: *gs://h01-release/data/20220326/c3/embeddings/segclr_csvzips/README* And precomputed annotations: <u>human</u>; <u>mouse</u>.

## Acknowledgements

We are grateful to Simon Kornblith, Geoffrey Hinton, Chen Sun, Dilip Krishnan, and Balaji Lakshminarayanan for helpful discussions. ALB, FC, CSSM and NMdC were supported by the Intelligence Advanced Research Projects Activity (IARPA) of the Department of Interior/Interior Business Center (DoI/IBC) through contract number D16PC00004; and by Allen Institute for Brain Science. The views and conclusions contained herein are those of the authors and should not be interpreted as representing the official policies or endorsements, either expressed or implied, of the funding sources including IARPA, DoI/IBC, or the U.S. Government. ALB, FC, CSSM and NMdC wish to thank the Allen Institute founder, Paul G. Allen, for his vision, encouragement, and support.

